# Dynamic Mode Decomposition for Transient Cavitation Bubbles Imaging in Pulsed High Intensity Focused Ultrasound Therapy

**DOI:** 10.1101/2024.02.26.582222

**Authors:** Minho Song, Oleg A. Sapozhnikov, Vera A. Khokhlova, Tatiana D. Khokhlova

## Abstract

Pulsed high-intensity focused ultrasound (pHIFU) can induce sparse *de novo* inertial cavitation without the introduction of exogenous contrast agents, promoting mild mechanical disruption in targeted tissue. Because the bubbles are small and rapidly dissolve after each HIFU pulse, mapping transient bubbles and obtaining real-time quantitative metrics correlated to tissue damage are challenging. Prior work introduced Bubble Doppler, an ultrafast power Doppler imaging method as a sensitive means to map cavitation bubbles. The main limitation of that method was its reliance on conventional wall filters used in Doppler imaging and optimized for imaging blood flow rather than transient scatterers. This study explores Bubble Doppler enhancement using dynamic mode decomposition (DMD) of a matrix created from a Doppler ensemble for mapping and extracting the characteristics of transient cavitation bubbles. DMD was first tested *in silico* with a numerical dataset mimicking the spatiotemporal characteristics of backscattered signal from tissue and bubbles. The performance of DMD filter was compared to other widely used Doppler wall filters - singular value decomposition (SVD) and infinite impulse response (IIR) highpass filter. DMD was then applied to an *ex vivo* tissue dataset where each HIFU pulse was immediately followed by a plane wave Doppler ensemble. *In silico* DMD outperformed SVD and IIR high pass filter and *ex vivo* provided physically interpretable images of the modes associated with bubbles and their corresponding temporal decay rates. These DMD modes can be trackable over the duration of pHIFU treatment using k-means clustering method, resulting in quantitative indicators of treatment progression.

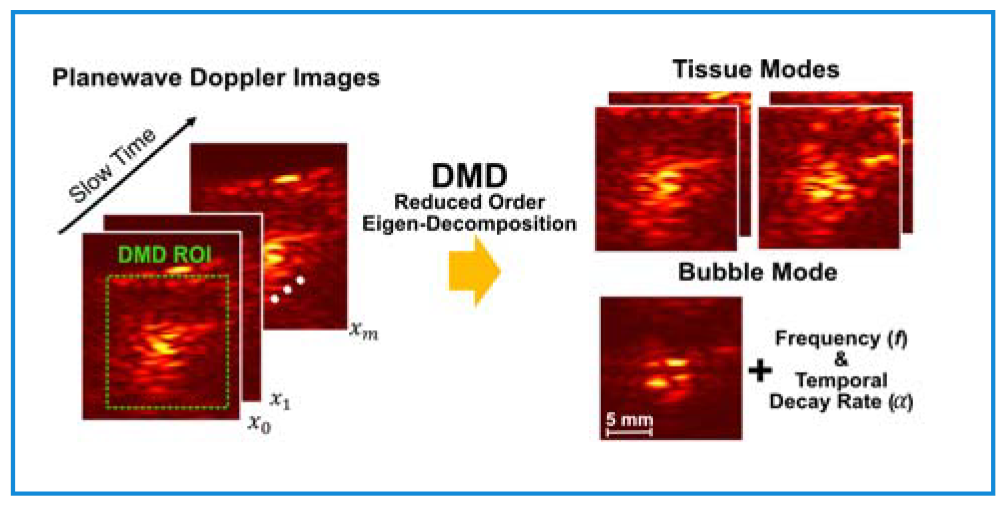

## I. Introduction

**P**ULSED high-intensity focused ultrasound (pHIFU) is a therapeutic ultrasound approach in which sparse *de novo* inertial cavitation is induced in tissue without the introduction of exogenous contrast agents, (*e*.*g*. microbubbles), resulting in mild mechanical disruption of the targeted tissue. pHIFU uses short (1-100 ms) pulses delivered at low duty cycle (<5%) and relatively low frequency (around 1 MHz) to avoid thermal effects and promote cavitation [1]–[8]. This regime was explored in conjunction with several clinical applications, including tumor permeabilization to enhance passive diffusion of systemically administered drugs [1], [2], [9], and inducing changes in tumor and benign tissue microenvironment [3]–[6]. The pHIFU treatments are commonly performed under real-time ultrasound (US) imaging guidance; conventional B-mode imaging with a probe integrated coaxially into the HIFU transducer is typically used for treatment planning and targeting. However, B-mode US has low sensitivity for detection of cavitation activity during treatment because of the transient nature of the bubbles and their rapid dissolution after HIFU pulse [7], [10]. Furthermore, even in cases where cavitation can be discerned in B-mode images as a hyperechoic region, this method does not provide a quantitative metric of cavitation or the resulting tissue disruption.

Conversely, passive methods, including passive cavitation detection (PCD) and passive acoustic mapping (PAM), quantify broadband acoustic emissions resulting from cavitation collapses (*i*.*e*. inertial cavitation) during HIFU pulses [7], [11]. These passive methods provide a sensitive quantitative metric of cavitation activity, but not of the resulting tissue damage, and the spatial information on cavitation occurrence is limited (*e*.*g*. axial resolution in PAM).

Bubble Doppler imaging is another active ultrasound method that was found to be more sensitive than B-mode and provide better spatial resolution than PAM in mapping bubbles induced by pHIFU [12]–[15]. In Bubble Doppler each HIFU pulse is immediately followed by an ensemble of planewave Doppler pulses. Those pulses encounter a distribution of residual cavitation bubbles that are rapidly (*i*.*e*. within milliseconds or the duration of the ensemble) changing due to dissolution. Doppler processing algorithm interprets those changes as motion, and thus the distribution of bubbles is displayed as an area of high Doppler power. The velocity associated with this area of high Doppler power is typically randomly fluctuating around zero, as the bubbles are not actually moving, only changing in size [12]. This creates a problem with selection of the wall filter used in conventional Doppler signal processing to remove clutter from stationary of slowly moving tissue: signal from nearly stationary yet temporally transient cavitation bubbles may be inadvertently filtered out. Furthermore, although Doppler power was found to correlate with PCD-derived broadband emissions, it does not provide any quantitative information on the cavitation bubble parameters, such as size or dissolution time, that could be linked to the level of tissue disruption.

One well-established wall filter is the infinite impulse response (IIR) high pass filter, which eliminates low frequency components from signals in the Doppler ensemble domain (*i*.*e*. slow time). This filter is effective when tissue motion is considerably slower than the motions of interest (*e*.*g*. blood flows speed vs surrounding tissue motion), and when prior knowledge about the appropriate cut-off frequency is available. However, the design of high pass filter and the cut-off frequency selection can be challenging in pHIFU-induced transient bubbles imaging due to the limited temporal sampling points during the bubbles’ lifespan [14]. Moreover, the quality of IIR filter can be degraded by transient clutter signal from HIFU reverberation.

Another commonly used wall filter in blood flow imaging, also proposed for cavitation imaging [14], [15], is singular value decomposition (SVD) filter [16]. The SVD filter decomposes a matrix of beamformed images, where each column is reshaped from an image corresponding to a Doppler ensemble pulse, into spatiotemporally coherent patterns (*i*.*e*. modes). These modes are orthogonal to one another and characterized by singular values that indicate the dominance of each mode. Filtering is achieved by reconstructing the image using a selective set of modes corresponding only to the tissues and motion of interest. The primary assumption underlying SVD filtering is that the amplitude of the clutter signal from background tissue consistently exceeds that from scatterers of interest, therefore, the modes corresponding to several largest singular values are removed. However, the amplitude of the backscattered signals from *de novo* cavitation bubbles can be comparable or even larger than those from tissue, making *a priori* mode selection challenging. Furthermore, because the residual bubbles typically dissolve within a few milliseconds after the HIFU pulse [12], a limited number of Doppler ensemble pulses can be acquired to capture the bubbles, resulting in degradation of the quality of SVD filter.

We propose the use of dynamic mode decomposition (DMD) as a technique to address the limitations of Bubble Doppler signal processing discussed above, namely – improve the reliability and sensitivity of detecting and mapping the cavitation bubbles and providing quantitative information on their parameters. DMD is a data-driven technique to analyze the dynamics of complex phenomena [17]. This technique originated in fluid dynamics to understand the coherent structures and patterns of high dimensional complex fluid flows [18], [19], but it then extended to various other fields including robotics control [20], imaging processing [21], neuroscience [22], plasma physics [23], and finance [24].

DMD decomposes time-series data into a reduced-order set of spatiotemporal modes, each of which consists of spatially coherent patterns and has the same behavior in time. While both SVD and DMD extract spatiotemporal information from the time-series snapshots, significant distinctions exist between the two. The spatial modes of SVD are not affected by the temporal order or arrangement of the input snapshots. In essence, SVD treats the input data more like a collection of images, focusing on its intrinsic structure and relationships between the individual images, rather than the temporal order in which the input data was collected. In contrast, DMD considers and captures both the intrinsic structure of the snapshots and their physically meaningful time behavior. Here, the DMD modes were assumed to represent quasi-harmonic motion with exponential growth or decay in amplitude. This is the most basic DMD implementation, but it matches the temporal behavior of Doppler signals backscattered from dissolving bubbles [12].

In this article, the feasibility of applying DMD processing for mapping the pHIFU-induced transient bubbles and identifying their temporal behavior was investigated. First, the performance of DMD was tested and compared to SVD and IIR high pass filtering in a numerically simulated image series that combined four different spatial patterns with varying time behaviors. Subsequently, DMD processing was applied to the planewave Doppler datasets acquired from pHIFU treatment of *ex vivo* tissue.

## II. Methods

### A. Experimental Arrangement and pHIFU Exposure Parameters

Fig. 1(a) shows the general schematic of the experimental setup that was used in the pHIFU exposures of *ex vivo* bovine tongue tissue. A 1.5 MHz HIFU transducer (8 cm aperture size, 6 cm focal distance and a 2 cm circular central opening) was controlled by high power driving electronics—a custom built 12-channel class D amplifier system [25], [26]. A 64-element ATL P7-4 ultrasound imaging probe was mounted at the center opening of the HIFU transducer and connected to Verasonics V1 system to acquire the planewave Doppler imaging data. The transducer assembly was mounted in a tank filled with degassed, deionized water. The *ex vivo* tissue sample was placed in front of the transducer in a holder attached to a 3D positioning system (Velmex Inc., Bloomfield, NY, USA), such that the HIFU focus was at a depth of 1 cm. The pHIFU pulsing protocol was the same as that previously used by our group for cavitation-based permeabilization of tissue [1], [12], [26]: pulse duration 1 ms, pulse repetition frequency (PRF) 1 Hz. During each HIFU pulse, the US imaging probe was operated in passive mode at a sampling rate of 20 MHz, to record broadband emissions0020from cavitation bubbles. The radio frequency (RF) signals were then bandpass and comb-filtered to remove backscattered HIFU harmonics and summed across the 64 channels with beamforming delays corresponding to the location of HIFU focus. The binary indicator of cavitation occurrence was obtained from the resulting PCD signal, and it was used to confirm the cavitation activity of the acquired dataset.

**Fig. 1.**
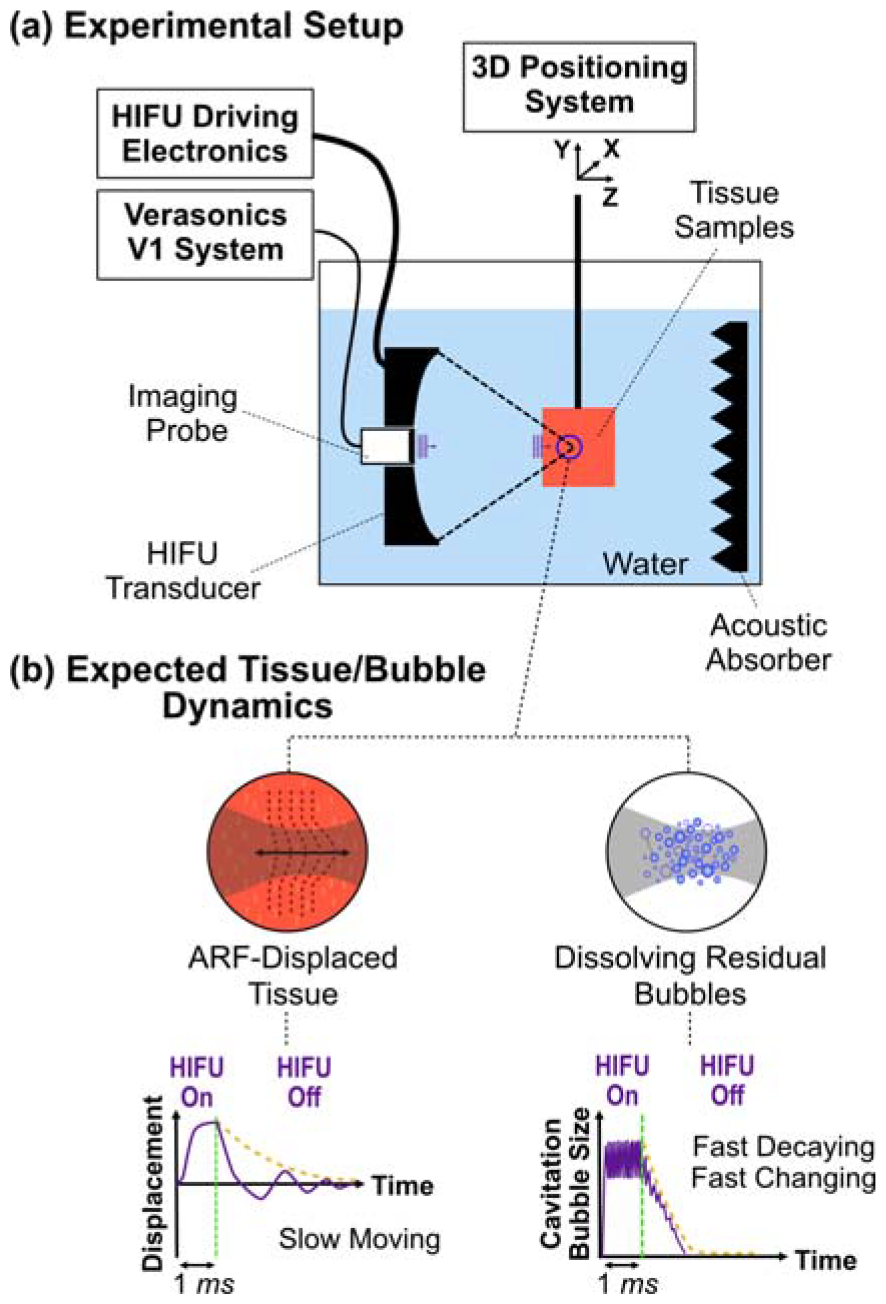
Schematic of (a) the experimental setup for pHIFU exposure in *ex vivo* bovine tongue tissue, (b) expected tissue and bubble dynamics. The elastic rebound motion of tissue displaced by acoustic radiation force was expected to be relatively slow with very gradual decay, whereas the distribution of residual cavitation bubbles of varying sizes was expected to rapidly change and dissolve.

Each HIFU pulse was immediately followed by a 5 MHz planewave Doppler ensemble consisting of 14 pulses at PRF of 3 kHz. Two different HIFU output levels were selected to achieve *in situ* focal pressure levels above and below the cavitation threshold. The *in situ* peak positive and negative pressures were calculated using hydrophone measurements in water reported previously [26] and a derating procedure for nonlinear waveforms [27] using attenuation coefficient of 0.044 Np/cm [28] with 1 cm depth in tissue, and were 20.9 / - 9.4 MPa, and 72.3 / -12.7 MPa, respectively. A total 30 and 60 HIFU treatment pulses were delivered to each focus location for non-cavitation and cavitation cases, respectively. The 4 focus locations were treated per output level, with similar results.

The bovine tongue tissue sample was obtained from a local abattoir and kept on ice until use within 48 hours. The choice of the bovine tongue was based on its anatomical structure, characterized by the interweaving of muscle bundles with fatty tissue, which presented a good cavitation probability [7]. The tissues were trimmed, cut fibrous surface, degassed in saline for 1 hour in a desiccant chamber, and embedded in low melting point agarose gel (UltraPure Agarose; Invitrogen) to facilitate its placement in the holder.

### B. Dynamic Mode Decomposition

#### 1) Underlying Assumptions

The target of DMD processing was to capture changes in the planewave Doppler imaging pulses backscattered from the HIFU focal area immediately after each HIFU pulse. The changes were expected to result from two phenomena: rapid dissolution of the cavitation bubbles of varying size and at varying rate, and slow elastic tissue rebound motion induced by acoustic radiation force (ARF) associated with the HIFU pulse, as shown in Fig. 1(b). We hypothesized that the captured changes in reconstructed images over slow time can be represented by a low-dimensional set of patterns, with each of these patterns associated with specific factors (*e*.*g*. cavitation or tissue motion) contributing to the observed changes. Additionally, linear approximation of the system dynamics was used as the simplest case, described by coupled linear differential equations:

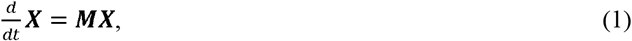

where ***X*** is a matrix of backscattered pulses that reflects the dynamics of tissue and bubbles, and ***M*** is a linear operator that revels the dynamics of the matrix ***X***. The linear approximation was expected to be adequate due to the limited number of temporal acquisition points (i.e. the number of pulses in the Doppler ensemble) within the bubbles’ lifespan.

#### 2) DMD Processing

Although DMD has been a prominent research field in recent years with many variations and advanced techniques developed, in this work we used the most basic implementation—exact DMD—suggested by Tu et al [29]. The data for pre-processing of DMD was constructed similarly to the general matrix formulation of SVD clutter filtering used in conventional Doppler imaging [16]. The input data were *m* +1 2D scan-converted in-phase and quadrature(I/Q) components acquired by Verasonics in slow time (labeled as *x*_0_,*x*_1_,…,*x*_*m*_), where *m* +1 is the number of pulses in Doppler ensemble. The total number of pixels in a single image (*x*_*i*_, *i* =0, …,*m*) was *n* with grid size corresponding to the one wavelength of Doppler ensemble pulse (∼0.3 mm). As illustrated in Fig. 2, each 2D I/Q dataset was reshaped into a column vector by attaching one column under the other in left-to-right order, and stacked into ***X*** and ***X***^′^matrices each having *m* columns and one time step shifted from one another(*i*.*e*. ***X*** contains *x*_0_ to *x*_*m*-1_ and contains ***X***^′^ *x*_1_ to *x*_*m*_). Then, the discrete time version of Eq. (1) could be written as

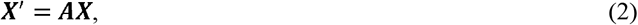

where ***A*** = *exp* (***M*** Δ *t*) and Δ *t* is time step in slow time. Notably, the number of rows (*n*) of the matrices ***X*** and ***X***^′^, corresponding to the total number of pixels in a planewave Doppler image, was on the order of 10^3^ in this study. It is usually much larger than the number of columns (*m*), which was on the order of 10.

**Fig. 2.**
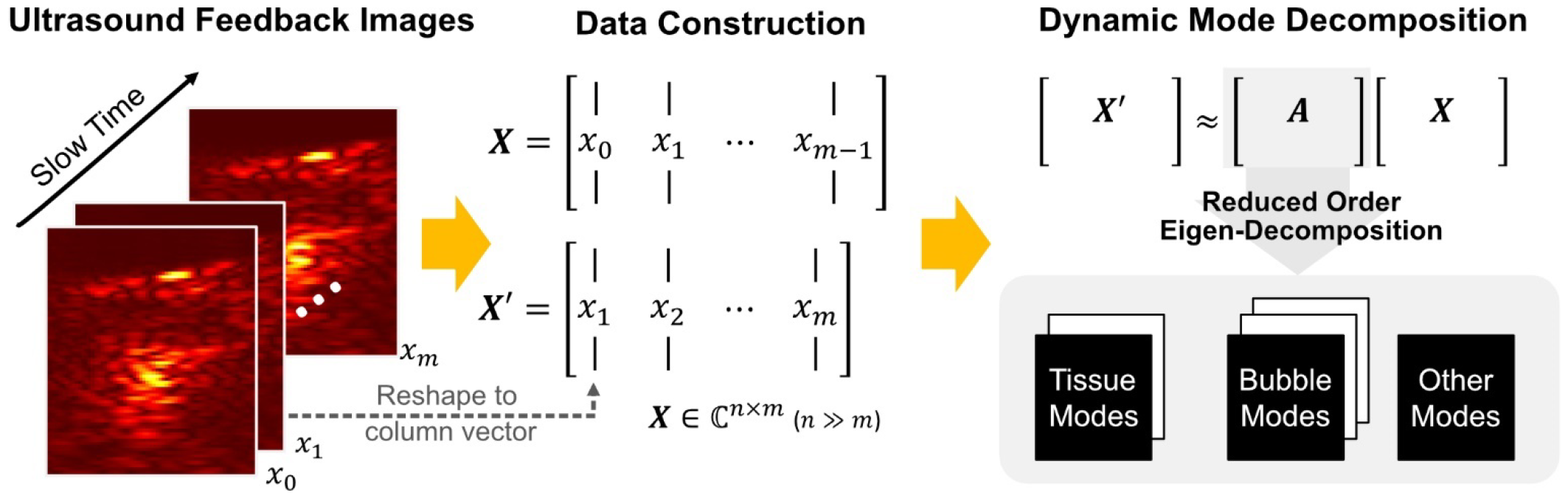
Schematic of the DMD processing. The scan-converted I/Q data were reshaped and stacked to form **X** and **X**^′^matrices with one time-step shift relative to each other. The DMD provides reduced ordered eigenvalues and DMD modes of ***A*** matrix which is the best-fitted operator satisfying **X**^′^≅ ***AX***. These modes can be physically interpretable as motion and/or changes in tissue or bubbles.

In (2), the matrix ***A*** fully describes the temporal evolution of the image dataset ***X***. However, the direct calculation of ***A***, as defined by

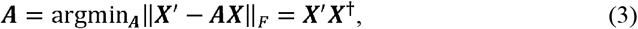

involves a computationally demanding process given that the dimensions of matrix ***A*** are *n* × *n*. In Eq. (3), denotes the process of determining the matrix ***A*** at which the argument attains its minimum, ‖ · ‖ _*F*_ is the Frobenius norm, and † indicates the pseudo-inverse. Instead of this direct computation, the exact DMD provides a reduced order set of exact eigenvalues and eigenvectors of matrix ***A***. Specifically, a rank-*r* approximation of eigen-decomposition of ***A***can be expressed as

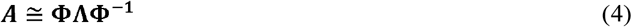

where **Φ**is an *n* × *r* matrix, its *r* columns of length *n* are eigenvectors of ***A*, Λ** is an *r* × *r* diagonal matrix in which each diagonal element is an eigenvalue of ***A***, and the superscript**-1**denotes the inverse of the given matrix. The objective of DMD is to derive matrices **Φ**and **Λ**. A comprehensive outline of the corresponding procedure is detailed in reference [29]; below is a brief summary:

a. Compute the SVD of the reconstructed dataset

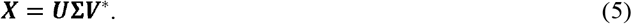

where the columns of ***U*** and ***V*** are right and left singular vectors with orthogonality that represent spatial modes and temporal coefficients, respectively, **Σ** is a diagonal matrix that has at most *m* non-zero singular values, and the superscript asterisk denotes the complex conjugate transpose.
b. Truncate the SVD matrices to rank-*r* by selecting the first *r* singular values in a descending order and corresponding columns of ***U*** and ***V***.(*r* ≤ *m*)Similar to SVD, the determination of the best-optimized rank *r* is a crucial step in DMD. For example, if the rank is too low, there is a risk of intertwining tissue and bubble modes, and if it is too high, meaningless clutter modes could complicate the interpretation of the modes and identifying modes associated with bubbles. In this study, *r* was determined when the summation of first *r* singular values exceeded 97% of that of all singular values (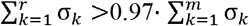, where σ_*k*_ is a k^th^ singular value or diagonal term of matrix **Σ**).

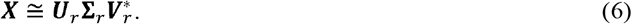

Where the subscript *r* indicates rank-*r* truncation of given matrix, such that the dimension of ***U***_*r*,_ **Σ**_*r*_, and 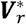 are *n* × *r,r* × *r* and *r* × *m*, respectively. Then, approximate ***A*** could be rewritten by substituting Eq. (6) into Eq. (3), as

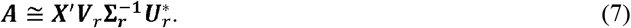
c. Compute the reduced order *r* × *r* matrix ***A***_***r***_, which is a similarity transformation matrix of ***A***.

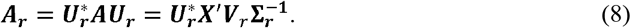
d. Compute the eigen-decomposition of ***A***_***r***_.

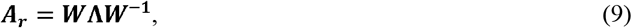

where ***W*** is the matrix whose columns are eigenvectors of ***A***_***r***_ and **Λ**is the diagonal matrix, whose diagonal elements are the eigenvalues of ***A***_***r***_. Since ***A***_***r***_ and ***A*** are related by matrix similarity, they share the same eigenvalues **Λ**, but different eigenvectors.
e. Compute eigenvector matrix **Φ** using Eqs, (4), (7), (8), and(9):

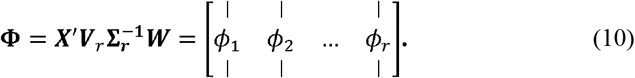

Where *ϕ*_*i*_ is i^th^ eigenvector (*i*.*e*. i^th^ column vector of **Φ**).

#### 3) DMD modes and eigenvalues

The diagonal elements of **Λ** and column vectors of **Φ** obtained from (9) and (10) represent the exact eigenvalue and corresponding eigenvector (DMD mode shapes) of matrix ***A***. The DMD mode shapes can be reshaped as 2D images, which are expected to correspond to physically interpretable modes such as planewave images of tissue and bubbles (Fig. 2). The continuous-time eigenvalue matrix **Ω**, defined as

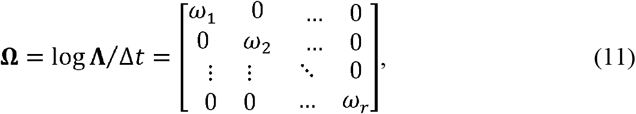

indicates how each DMD mode shape evolves in time. Here, ω_*i*_ is i^th^ continuous-time eigenvalue (*i*.*e*. i^th^ diagonal element of **Ω**). The negative real part of ω_*i*_ corresponds to the temporal decay rate; the imaginary part corresponds to the angular frequency or rate of change.

#### 4) Mode Classification and Reconstruction

Over the course of pHIFU treatment, the behavior of the transient bubbles at the focus is expected to change as the tissue becomes more disrupted. To identify and monitor these changes, the discrete-time eigenvalues (i.e. diagonal elements of **Λ**) were classified and tracked on the complex plane using k-means clustering method, which is a widely used method that groups dataset into k clusters by minimizing the distance between each data point and its cluster’s centroid. Since the discrete-time eigenvalues for exponentially decaying dynamics are bounded by a unit circle on the complex plane, they are more suitable for clustering than continuous-time eigenvalues (ω_*i*_). The classified eigenvalues and corresponding DMD mode shapes were then characterized with respect to the three metrics: frequency, temporal decay rate, and maximum power level of the DMD mode shape defined as *p*_*max*_ = *max* (∣ ϕ ∣^2^). Based on the metrics, each DMD mode could be associated with the dynamics of tissue or bubbles, and selective reconstruction of those modes could be performed. A bubble mode reconstruction, for example, can be expressed as

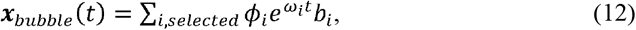

where *i, selected* indicates the index of selected modes that were interpreted as components of bubble mode and *b*_*i*_ is i^th^ element of initial mode amplitude vector ***b*** related to the initial image *x*_0_ (*i*.*e*. ***b*** *=* **Φ** ^†^ *x*_0_).

#### 5) Connection of DMD Modes to Conventional Doppler Variables

Elimination of certain DMD modes can serve as a wall filter in conventional Doppler processing. For each DMD bubble mode, the consecutive bubble images in (12) (e.g. ***x*** _*bubble*_ at *t*_1_= Δ*t* and *t*_2_= 2 Δ*t*) have a phase shift of *Im* { ω} Δ*t* relative to each other over all the image pixels, where *Im* { ω}denotes the imaginary part of ω. This means that DMD processing directly provides the phase shift necessary for color Doppler estimation of velocity associated with each mode. Specifically, the angular frequency of a DMD mode (*i*.*e. Im* { ω}) corresponds to Doppler phase shift, from which Doppler velocity is calculated as:

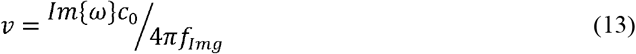

where *c*_0_ is the sound speed in tissue and *f*_*Img*_ indicates the frequency of planewave Doppler imaging pulses. This also implies that a positive DMD frequency corresponds to motion directed towards the HIFU transducer, whereas a negative frequency corresponds to motion away from it.

### C. Evaluation of DMD performance in a Numerical Example

The performance of DMD was evaluated in a numerically simulated time series of images, meant to represent a simplistic motion from tissue and bubbles with PRF and number of pulses matching the experimental arrangement. The data were processed with DMD and by commonly used wall filters in conventional Doppler processing – SVD and IIR high pass filter – for side-by-side comparison. The similar comparison study between DMD and other modal decomposition methods – principal component analysis (PCA) and independent components analysis (ICA) – can be found in neuroscience paper [22]. As shown in Fig. 3(a), the data was a combination of four shapes ***S***_1,_ ***S***_2_, ***S***_3_and ***S***_4_: a square of uniform brightness, two circles with radially cosine distribution of brightness, and random noise, respectively. Each of them except random noise ***S***_4_ was spatiotemporally coherent – brightness of all points of the shape had the same behavior in time. The input brightness data ***X***[*n*] was thus defined as

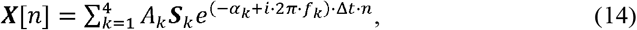

where *A*_*k*_ was the initial brightness amplitude of each shape, *f*_*k*_ was frequency (*f*_1_ = 50 Hz, *f*_2_= 500 Hz, *f*_3_= 550 Hz, and *f*_4_= 0 Hz), α_*k*_ was temporal decay rate (α_1_ = 0.1 *ms*^-1^, α_2_ = 2 *ms*^-1^, α_3_ = 5 *ms*^-1^, and α_4_ = 0 *ms*^-1^), Δ*t* was time step Δ*t* = 0.33*ms*, which corresponded to the experimentally allowable PRF of 3 kHz), and *n* was sample number or the virtual imaging pulse number (1≤ *n* ≤14). At each time step, random noise ***S***_4_ was repeatedly generated to ensure that there was no spatial or temporal coherence. The initial brightness amplitudes, frequencies, and temporal decay rates of the modes were chosen to approximate the ultrasound images of slowly moving tissue which typically dominates the image amplitude, and that of a collection of bubbles which are quickly changing and dissolving. The initial brightness amplitudes of each shape (*A*_1_, *A*_2_, *A*_3_, and *A*_4_) were 2, 1, 1, and 0.01, respectively. For tissue containing two bubble areas (*A*_2_and *A*_3_), the amplitudes were designed to have signal-to-noise ratio (SNR) of 40 dB vs random noise level (*A*_4_), and that of tissue (*A*_1_) was set to be twice larger than *A*_2_ or *A*_3_ Figure 3(b) shows the SNR changes of each mode over slow time. The brightness of the two area of bubbles (***S***_2_ and ***S***_3_) rapidly decayed to below the random noise level within a few sampling points (3 sampling points (∼ 1 *ms*) and 6 sampling points (∼2.2 *ms*), respectively), which corresponded to a previously reported decay time for residual cavitation bubbles [12].

**Fig. 3.**
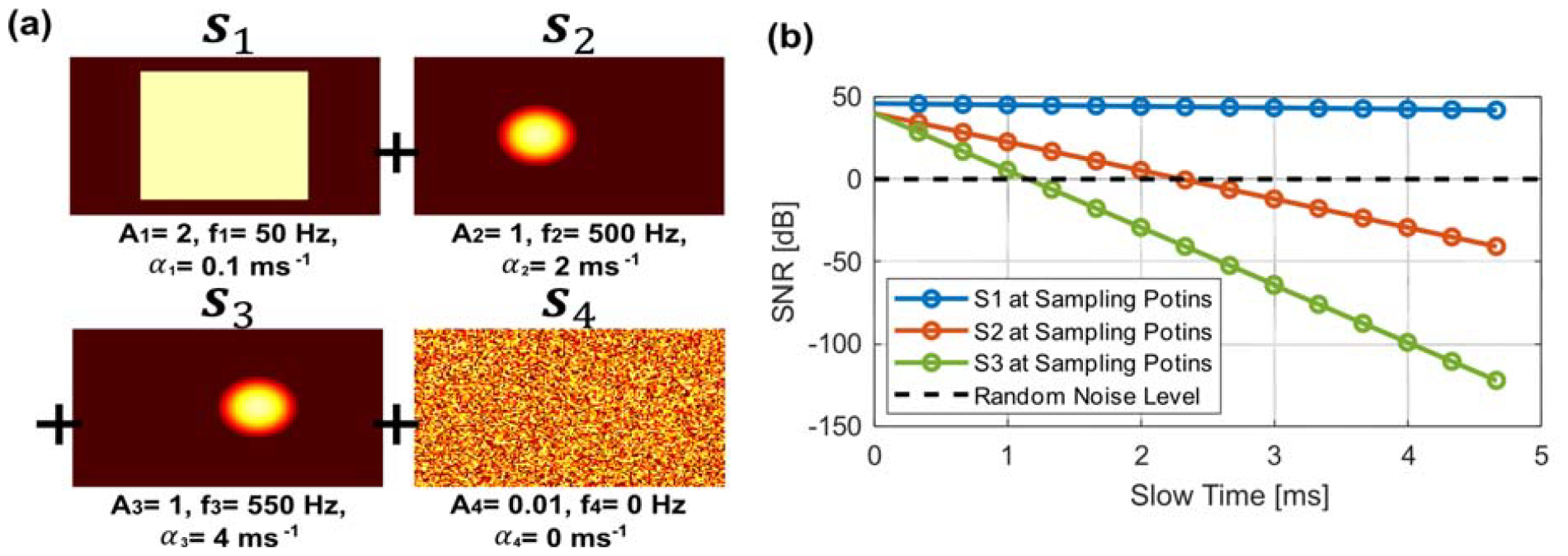
Numerical example for testing DMD performance vs SVD and IIR high pass filter. (a) The input data consisting of four components of simulated 2D scan data: a square image with high amplitude, low frequency, and low decay rate, simulating tissue backscatter (S_1_), left and right-side circles inside the square with low amplitude, high frequency, and high decay rate (S_2_, S_3_), simulating backscatter from bubbles, and a low amplitude random noise without any coherence in space and time (S_4_). (b) Illustration of the amplitude decay for each component of input data across slow time. The amplitude was expressed in SNR, as compared to the random noise level.

Three different methods were applied to the input data described above: DMD, SVD, and IIR high pass filter. In SVD, the data processing followed the same procedure as that described for DMD until step b). IIR high pass filter with the cut-off frequency of 275 Hz and order 3 was utilized to eliminate the image of background tissue (*i*.*e*., ***S***_1_). To assess the robustness of the DMD from random noise ***S***_4_, the simulation was repeated n=1000 times with random noise regenerated every time. Mean and standard deviation of the frequency and temporal decay rate for each DMD mode, as measured by each of the processing methods, were obtained and compared to ground truth.

## III. Results

### A. Evaluation of DMD performance in the Numerical Example

Fig. 4(a), (b), and (c) display representative cases of the numerical dataset processed with the three methods: four DMD modes along with their corresponding frequency (*f*) and temporal decay rate (α), the first four columns of SVD ***U*** matrix with the corresponding singular values (σ), and the result of IIR high pass filter, respectively. For DMD, each mode image, and corresponding values of *f* and αwere very consistent across repeated simulation with regenerated random noise. Table I shows the mean and standard deviation of *f* and αestimated by DMD. As seen, DMD allowed to effectively decompose spatially coherent patterns and provided *f* and α estimates with high accuracy, within 0.6% error vs ground truth. The modes 1, 2, and 3 closely corresponded to the shapes of input data without overlapping with each other. However, mode 3 exhibited more contamination by the random noise compared to modes 1 or 2. This was expected, as the amplitude of mode 3 brightness exceeded that of random noise for only 3 out of 14 sampling points. In contrast, SVD was not able to separate modes 2 and 3 (“bubble modes”), and, as seen in Fig 4(b), they overlapped. The singular value of mode 1 (“tissue mode”) was an order of magnitude higher than those of modes 2 and 3, which is consistent with the input dataset where mode 1 had the highest initial amplitude and the lowest α. Fig. 4(c) shows the result of applying IIR high pass filter to the data, meant to eliminate mode 1, leaving only modes 2 and 3, and the result of subtraction of the filtered data from the input data, meant to reconstruct mode 1. Because the appropriate cut-off frequency to filter out mode 1 was known a priori, the separation of the “bubble modes” from “tissue mode” was successful, and the images of the modes had minimal overlap.

**TABLE I.** The Statistical Results of Frequency and Temporal Decay Rate Estimated by DMD (N=1000)

|  |  | Mode 1 | Mode 2 | Mode 3 |
| --- | --- | --- | --- | --- |
| $f$<br>[Hz] | Ground truth | 50 | 500 | 550 |
|  | Mean | 49.980 | 500.293 | 553.160 |
|  | Std | 0.007 | 0.763 | 3.431 |
| $\alpha$<br>[ms<sup>-1</sup>] | Ground truth | 0.1 | 2 | 4 |
|  | Mean | 0.1008 | 2.0003 | 4.0093 |
|  | Std | 0.0001 | 0.0034 | 0.0189 |

**Fig. 4.**
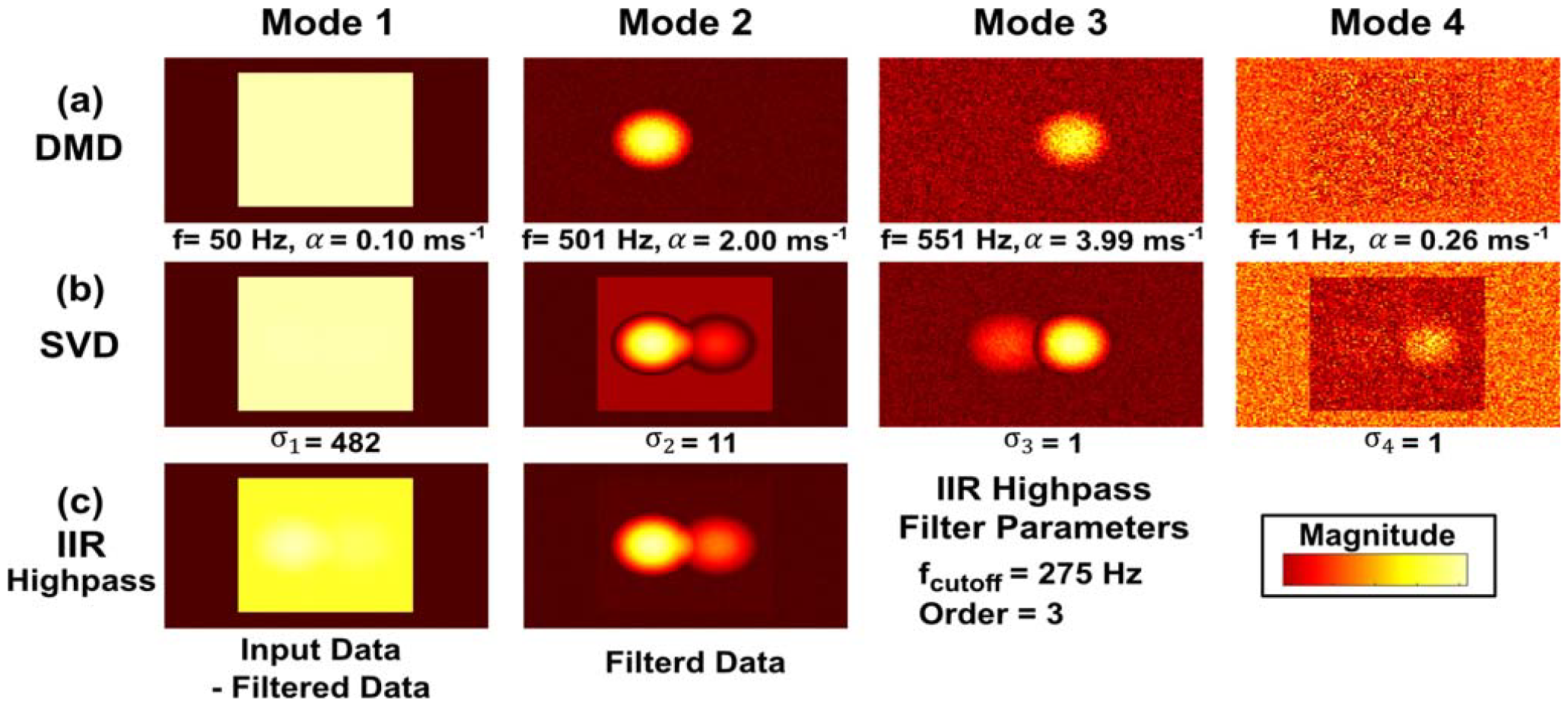
Separation of modes from numerical example with the different filtering methods: DMD, SVD, and IIR highpass. (a) Four DMD modes and their corresponding frequencies and temporal decay rate. (c) Four SVD modes corresponding to the first four columns of SVD ***U*** matrix with corresponding singular values σ. (d) IIR high pass filtered data and its subtraction from original data.

### B. DMD of Doppler Data for ex vivo pHIFU Exposures without Cavitation

During the first set of pHIFU exposures in *ex vivo* bovine tongue tissue, the pHIFU amplitude was set to be below the cavitation threshold, as confirmed by PCD. A representative example of DMD modes corresponding to such a HIFU pulse, with corresponding *f* and α are shown in Fig. 5. The DMD-processed region of interest (ROI) is shown within the larger B-mode image, and the pHIFU beam location is denoted by a green dashed V-shape line on each DMD mode image. All DMD modes were normalized to the maximum amplitude across all identified modes and were displayed with adjusted range of color bar for better visualization. Three DMD modes were identified, all with frequencies below 0.1 kHz. The first mode showed nearly stationary tissue speckles with the highest amplitude. The second and third modes, characterized by significantly lower amplitude levels (∼15% of the first), showed a paired behavior. The mode images, temporal decay rates, amplitude levels, and absolute values of frequency were very similar, and the only difference was the opposite sign of the frequency, indicating the opposite directions of axial motion of the two modes. The Doppler velocity corresponding to the last two modes, calculated from (13), was 1.4 - 1.5 cm/s. These values align with the expectations based on ARF-induced elastic rebound of tissue, as shown in our previous study [30].

**Fig. 5.**
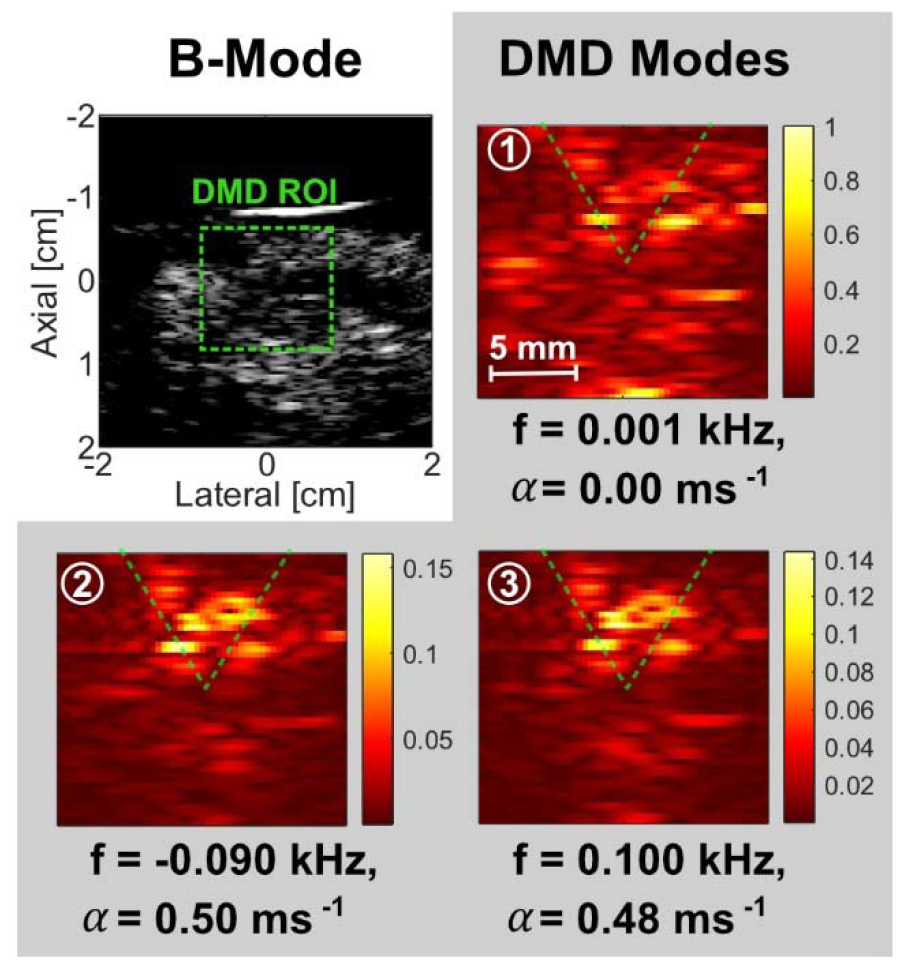
DMD modes of bubble Doppler obtained from pHIFU exposure of *ex vivo* tissue without cavitation. The green box on the B-mode image highlights the region of interest for bubble Doppler. Frequencies and decay rates are presented at the bottom of the corresponding mode images. The color bar scale was normalized to the maximum amplitude value over all DMD modes. HIFU was delivered from top of the images to the target point indicated by V-shaped green dashed line.

The k-means clustering results for discrete-time eigenvalues (**Λ**) over 30 HIFU pulses on the complex plane are shown in Fig. 6(a). Each eigenvalue was denoted by a colored cross, and each identified cluster was assigned a different color. Three clusters were identified, each of which showed tight clustering with maximum eigenvalue distances of 0.041, 0.016, and 0.014, respectively. Fig 6(b) and (c) show the values of *f* and αover 30 HIFU pulses; as seen, those values remained nearly constant, as expected for pHIFU exposures below the cavitation threshold.

**Fig. 6.**
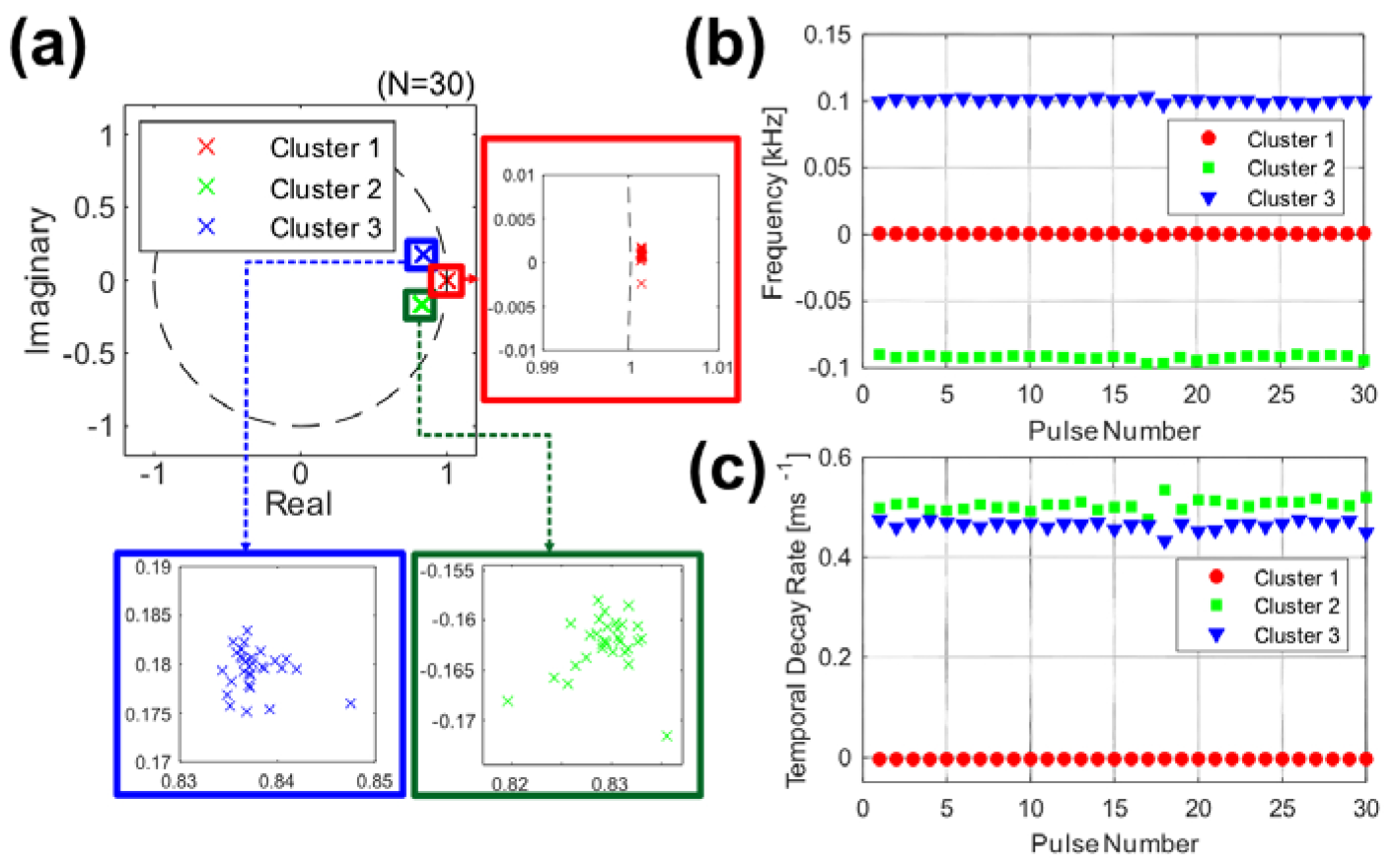
DMD eigenvalues during the 30-pulse pHIFU exposure without cavitation. (a) The discrete time eigenvalues for all 30 pHIFU pulses on the complex plane and the results of their k-means clustering. Three eigenvalue clusters were identified and marked with different colors. The (b) frequency and (c) temporal decay rate of the three clusters plotted vs HIFU pulse number.

### C. DMD of Doppler Data for ex vivo pHIFU Exposures Inducing Cavitation

A representative example of DMD modes corresponding to a HIFU pulse that induced cavitation in the *ex vivo* tissue is shown in Fig. 7, in the same format as Fig. 5. DMD processing consistently yielded five modes for those pHIFU exposures that represented the dynamics of tissue and residual bubbles immediately following each pulse. The first DMD mode displayed speckles over the entire ROI, with brighter hyperechoic regions at the focus. This mode had the lowest frequency (0.006 kHz) and temporal decay rate (0.01 *ms*^-1^), indicating stationary backscatter from tissue and persistent residual bubbles that did not change within the imaging time window. The second mode was similar to the first mode in speckle distribution but had a higher temporal decay rate. The third and fourth modes depicted the speckles around the focus, with higher frequencies and temporal decay rates than the first two modes and lower amplitude level (<40% of the second). The Doppler speeds corresponding to those two modes were 2.4 and 3.7 cm/s with opposite directions, likely linked with ARF-displaced tissue motion similarly to second and third modes in Fig 5. The fifth mode showed a backscatter pattern highly localized to the focal region, and the corresponding frequency, temporal decay rate, and Doppler speed were the highest among all modes (0.785 kHz, 3.23 *ms*^-1^, and 12.1 cm/s). This mode was thus hypothesized to be associated with quickly dissolving cavitation bubbles. Indeed, previous studies have shown that most cavitation bubbles in pHIFU exposures dissolved within 1-2 *ms* following the pulse, which is consistent with of this mode. The amplitude ratio between the first and fifth mode was initially -2 dB but reached -30 dB after 1 *ms*. Further, many cavitation bubbles with different radius and dissolution rate are expected to be present within the resolution cell of planewave image. Bubbles dissolving at different rates may be interpreted by DMD as rapidly changing backscattered signal.

**Fig. 7.**
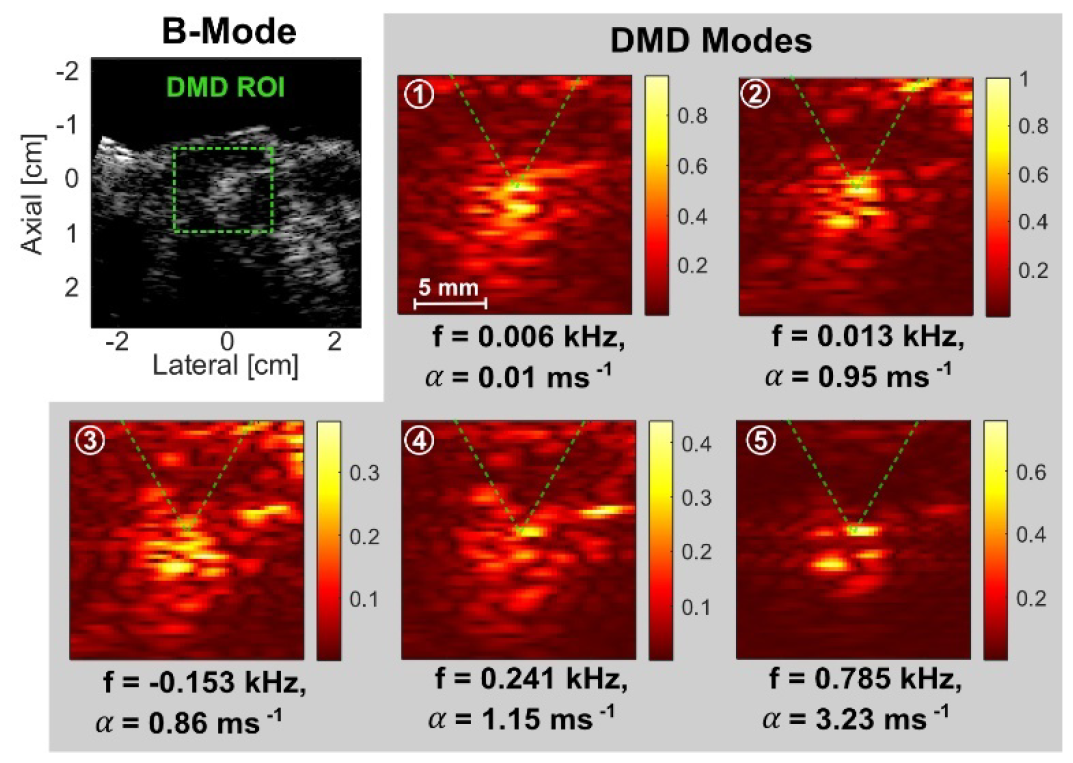
DMD modes of Bubble Doppler obtained from pHIFU exposure of *ex vivo* tissue with cavitation. The green box on the B-mode image highlights the region of interest for bubble Doppler. Five DMD modes were identified, and the corresponding images are presented with frequencies and decay rates. The color bar scale was normalized to the maximum amplitude value over all DMD modes. HIFU was delivered from the top of the images to the target point indicated by V-shaped green dashed line.

The evolution of the DMD eigenvalues and modes over the 60 pHIFU pulses delivered to the same location of the tissue sample is presented in Fig. 8. The results of clustering the discrete time eigenvalues over pHIFU pulses on the complex plane are shown in Fig. 8(a) in the same format as Fig. 6(a). Five clusters were identified corresponding to the five DMD modes in Fig. 7 and labeled with different colors. In particular, cluster 5 corresponding to mode 5, likely to be representative of cavitation, was assigned purple color. The eigenvalues for cluster 5 were clustered more loosely compared to the others; the maximum distances of eigenvalues in each cluster were 0.017, 0.221, 0.198, 0.419, and 0.521, respectively.

**Fig. 8.**
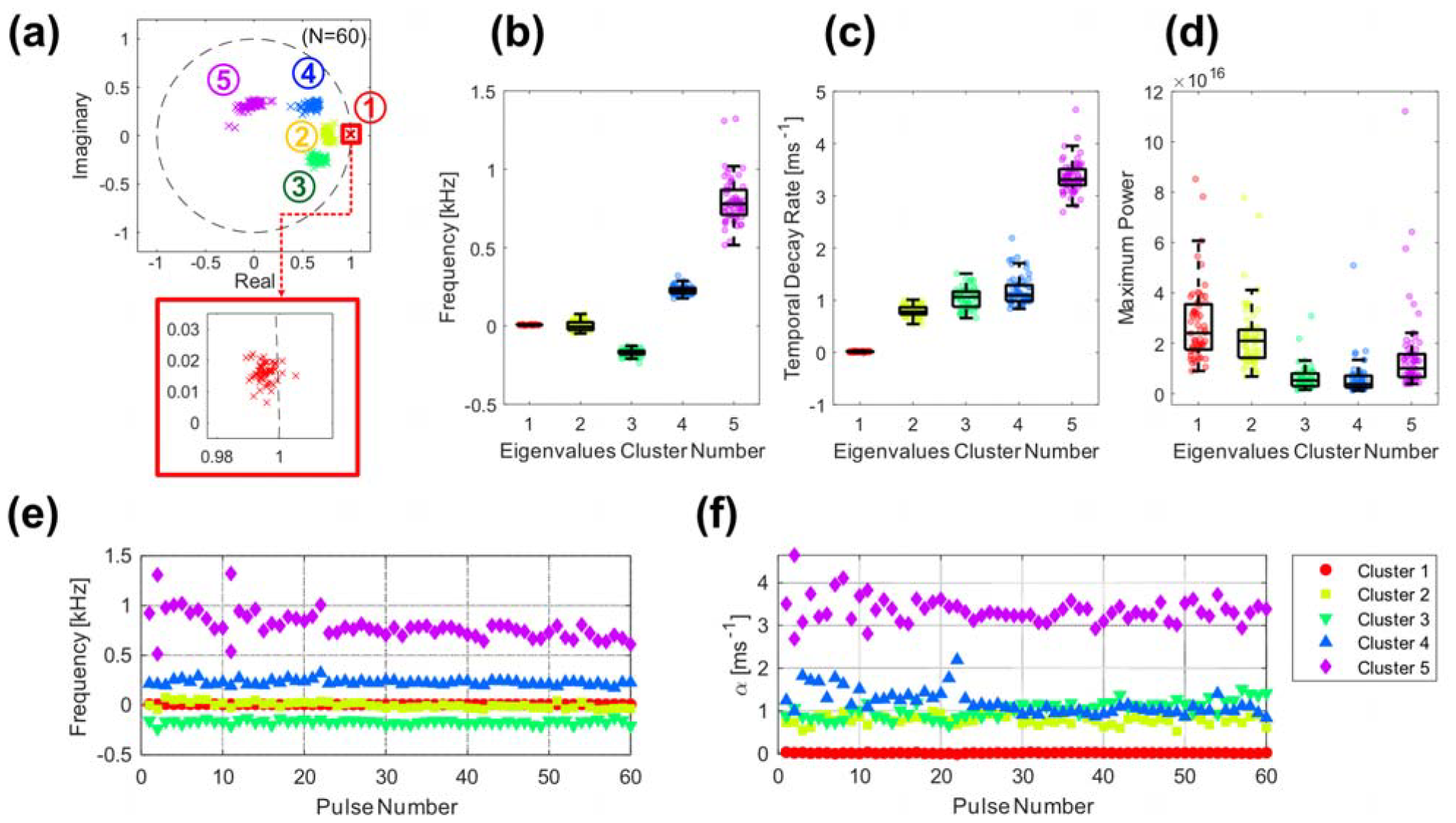
DMD eigenvalues during the pHIFU exposure with cavitation. (a) The discrete time eigenvalues over 60 pHIFU pulses on the complex plane and the results of their k-means clustering. Five eigenvalue clusters were marked as different colors and numbers. The box plot for (b) frequency, (c) temporal decay rate, and (d) maximum power of each eigenvalue cluster over all pulses; error bars represent minimum and maximum data points within 1.5 times the interquartile range (IQR), defined as the distance between the 0.25 and 0.75 quantile (also termed whisker). The (e) frequency and (f) temporal decay rate plotted vs HIFU pulse number.

Fig. 8(b), (c), and (d) show the box plots for the values of frequency, temporal decay rate, and maximum power for each eigenvalue cluster, for the 60 HIFU pulses. Clusters 1 and 2 showed closely distributed *f* and α, while the maximum power distribution was broad. This reflects the fact that the hyperechoic region at the focus attributed to the persistent residual bubbles increased in brightness over the course of the exposure. On the other hand, cluster 5 exhibited a broad range of *f* from 0.520 kHz to 1.320 kHz and high αwith a median of 3.3*ms*^-1^, supporting the hypothesis that it represented the rapidly dissolving residual cavitation bubbles. Additionally, clusters 3 and 4 showed tighter distribution ranges in *f* and α than cluster 5 and were characterized by the smallest maximum power values.

Fig. 8(e) and (f) show the evolution of *f* and αover the 60 HIFU pulses. The frequency for clusters 1 to 4 and temporal decay rate for clusters 1 to 3 were almost constant, similarly to clusters 1-3 in the exposure without cavitation (Fig. 6(b) and (c)). Conversely, in cluster 5, the frequency gradually decreased from around 1 kHz to 0.6 kHz as the treatment progressed. The temporal decay rate in clusters 4 and 5 decreased slightly (*i*.*e*. became less decaying), from around 2 to 1.5 *ms*^-1^ and 4 to 3*ms*^-1^, respectively, and became less variable after about 20^th^ pulse. Taken together, those changes suggest that the distribution of bubble sizes, positions and decay rates became more consistent in the second half of pHIFU exposure, which could indicate consolidation of the bubbles in the same disrupted areas of tissue and not inflicting more damage.

## IV. Discussion and Conclusions

This study investigated the utility of DMD technique in imaging transient cavitation bubbles during pHIFU therapy. The effectiveness of DMD to identify spatially coherent patterns and quantify their temporal dynamics was first demonstrated in a numerically simulated image dataset and compared to the performance of SVD and IIR wall filtering. Then DMD was applied to planewave Doppler datasets acquired during pHIFU exposures of *ex vivo* tissue.

In the simulation study DMD outperformed the SVD and IIR filter in terms of quality of pattern decomposition and accuracy of the temporal dynamics metrics associated with them–frequency (*f*) and temporal decay rate (α). The values of *f* and α obtained through DMD were remarkably accurate for all modes, despite the simulation’s input data containing two limitations that are inherent in the experimental implementation. First, the total duration of Doppler ensemble in slow time (*i*.*e*. 4.7 for 14 sampling points with a 3 kHz PRF) was insufficient to encompass even a single cycle of 50 Hz, which was designated as the frequency for the square shape. Second, the SNR for the two circles were intentionally reduced to levels below random noise after 6 and 3 sampling points, respectively, as shown in Fig. 3(b).

In pHIFU exposures of *ex vivo* tissue, DMD modes and their *f* (or corresponding Doppler speed) and *α* were found to be physically interpretable and consistent with prior studies. In pHIFU exposures without cavitation (Fig. 5), the three DMD modes could be identified as stationary tissue (first mode) and ARF-displaced axial tissue motion in both directions (second and third modes). This is because the tissue, immediately after the termination of HIFU pulse, is expected to move towards the HIFU transducer, and then to rebound away within the total acquisition duration (*i*.*e*. 4.7 *ms*), as seen prior study [30]. Similarly, each DMD mode in pHIFU exposures resulting in cavitation (Fig 7) can be identified as stationary tissue (first and second modes), ARF-induced axial tissue motion in both directions (third and fourth modes), and transient bubbles (fifth mode). Comparison with the non-cavitation modes from Fig 5 showed that the modes corresponding to the stationary tissue and ARF-induced tissue motion in Fig. 7 had a bright area at the focus, suggesting a contribution from residual bubbles. This may be attributed to the fact that some larger size, more stable bubbles moved together with tissue and potentially amplified the ARF displacement [30].

In the bubble mode, the temporal decay rate α can be interpreted as an average rate of bubble dissolution within a resolution cell of a planewave image. However, the physical interpretation of frequency *f*, or the corresponding Doppler speed, is not as straightforward, considering that residual bubbles are not expected to move or oscillate after the HIFU pulse. A possible explanation is that the speed of the surface of shrinking bubbles within the resolution cell is synchronized, leading to a phase shift in slow time. The DMD then interprets this as motion.

The traceability of tissue and bubble modes is another advantage of DMD. As seen in Fig. 8, the k-means clustering on complex planes allows one to track each DMD mode. This ability is particularly useful for quantitative imaging guidance in pHIFU therapy, as it allows for tracking of bubble characteristics, which could be physically interpretable in terms of inflicted tissue damage. For example, the decrease in frequency and temporal decay rate for cavitation bubble mode, observed in Fig 8(e) and (f), could be interpreted as a sign of bubble coalescence and decrease in surrounding tissue stiffness.

In this study the utility of DMD was explored for a relatively narrow application – as a type of wall filter in Doppler-based detection, mapping and quantitation of cavitation bubbles induced by pHIFU. However, we anticipate this technique to be useful in a number of other time-resolved ultrasound imaging tasks, e.g. vascular imaging, characterization of flow patterns in large veins and arteries, and monitoring of tissue ablative interventions over time.

This study is not free of limitations that warrant further investigation. First, the clustering method meant to track the different modes over the course of pHIFU exposure did not include automatic identification of the cluster corresponding to the cavitation bubbles. To automate the labeling process, several algorithms could be considered, including the eigenvalue thresholding (using *f* and α), mode image-based metrics (*e*.*g*. contrast-to-noise ratio), or machine learning techniques. Second, the appropriate *r* selection for rank-*r* truncation using SVD, described in step b) of DMD processing section, also affects the decomposition quality of DMD. As mentioned in the methods, a preliminary empirical investigation was performed to identify the most effective truncation method for fast-dissolving bubbles. Such an investigation may need to be repeated if there are changes in acquisition parameters, such as the number of imaging pulses (*m*) used and Doppler PRF. The selection problem in SVD is well known and previously reported in works on Doppler clutter filtering [14], [15]. Most importantly, current study only involved pHIFU exposures of stationary *ex vivo* tissue, devoid of circulation or physiological motion. On the one hand, application in the *in vivo* environment will undoubtedly complicate the analysis and increase the number of modes; on the other hand, the advantages of DMD are expected to stand out more. This will be explored in future studies.

## Notes

This work was supported by NIH under Grant R01EB023910, and 2R01CA154451.

### Competing Interest Statement

The authors have declared no competing interest.

